# Genetic mechanisms of primary chemotherapy resistance in pediatric acute myeloid leukemia: A report from the TARGET initiative

**DOI:** 10.1101/475376

**Authors:** Nicole A. McNeer, John Philip, Heather Geiger, Rhonda E. Ries, Vincent-Philippe Lavallée, Michael Walsh, Minita Shah, Kanika Arora, Anne-Katrin Emde, Nicolas Robine, Todd A Alonzo, E. Anders Kolb, Alan S Gamis, Malcolm Smith, Daniela Se Gerhard, Jaime Guidry-Auvil, Soheil Meshinchi, Alex Kentsis

**Affiliations:** Department of Pediatrics, Memorial Sloan Kettering Cancer Center, New York, NY; Fred Hutchinson Cancer Research Center, Seattle, WA; Computational and Systems Biology Program, Sloan Kettering Institute, Memorial Sloan Kettering Cancer Center, New York, NY; New York Genome Center, New York, NY; Department of Biostatistics, University of Southern California, Los Angeles, CA; Nemours Center for Cancer and Blood Disorders, Nemours/Alfred Dupont Hospital for Children, Wilmington, DE; University of Missouri-Kansas City School of Medicine, Kansas City, MO; National Cancer Institute, Rockville, MD; Molecular Pharmacology Program, Sloan Kettering Institute, Memorial Sloan Kettering Cancer Center, New York, NY; Department of Pediatrics, Pharmacology, and Physiology & Biophysics, Weill Medical College of Cornell University, New York, NY

## Abstract

Acute myeloid leukemias (AML) are characterized by mutations of tumor suppressor and oncogenes, involving distinct genes in adults and children. While certain mutations have been associated with the increased risk of AML relapse, the genomic landscape of primary chemotherapy resistant AML is not well defined. As part of the TARGET initiative, we performed whole-genome DNA and transcriptome (RNA and miRNA) sequencing analysis of pediatric AML with failure of induction chemotherapy. We identified at least three genetic groups of patients with induction failure, including those with *NUP98* rearrangements, somatic mutations of *WT1* in the absence of *NUP98* mutations, and additional recurrent variants including those in *KMT2C* and *MLLT10.* Comparison of specimens before and after chemotherapy revealed distinct and invariant gene expression programs. While exhibiting overt therapy resistance, these leukemias nonetheless showed diverse forms of clonal evolution upon chemotherapy exposure. This included selection for mutant alleles of *FRMD8*, *DHX32*, *PIK3R1*, *SHANK3*, *MKLN1*, as well as persistence of *WT1* and *TP53* mutant clones, and elimination or contraction of *FLT3*, *PTPN11*, and *NRAS* mutant clones. These findings delineate genetic mechanisms of primary chemotherapy resistance in pediatric AML, which should inform improved approaches for its diagnosis and therapy.

## Introduction

Overall survival for children with acute myeloid leukemia (AML) remains low, due principally to the failure to achieve durable disease remission after initial induction therapy. Failure rate of primary induction remission therapy in pediatric AML is 10-15%, and only about a third of patients for whom primary induction therapy fails are ultimately cured (1). Reasons for the lack of response to initial chemotherapy in pediatric AML remain unclear, and a molecular understanding of this process is needed. Since the first AML genome was sequenced (2, 3), numerous genomic profiling studies have revealed diverse disease subtypes and distinct genetic modes of disease relapse (4). For example, whole-genome sequencing of AML specimens from adults with relapsed disease revealed broad patterns of clonal evolution, suggesting that either founding clones gained mutations upon relapse, or that diagnostic subclones persisted with acquisition of additional mutations after therapy (5). Analysis of whole exome capture sequencing from matched diagnosis, remission, and relapse trios from twenty pediatric AML cases showed that responses of specific genetic clones were associated with disease relapse (6). Similarly, clonal persistence after induction chemotherapy was found to be associated with disease relapse in adult AML (7).

Recent study of primary chemotherapy resistance in a cohort of 107 children and adults with AML using targeted gene sequencing demonstrated that few patients exhibited specific individual mutations associated with primary chemotherapy resistance and failure of induction chemotherapy (8). In addition, at least for some patients, chemotherapy resistance is caused by the epigenetic activation of the transcription factor MEF2C (9-10). This suggests that there are additional genetic or molecular mechanisms mediating primary chemotherapy resistance in pediatric and adult AML. Importantly, pediatric AML is characterized by distinct genetic mutations and genomic rearrangements, with relative paucity of the recurrent mutations frequently observed in adult AML (11). Thus, direct study of primary chemotherapy resistance in pediatric AML is needed.

Here, we assembled a cohort of pediatric patients with primary chemotherapy resistance and failure of induction chemotherapy, as part of the TARGET AML initiative. We analyzed whole-genome DNA, mRNA, and miRNA sequence data, obtained at diagnosis and upon chemotherapy administration. These studies revealed distinct classes of genetic mutations and their clonal evolution in chemotherapy resistant disease, which should inform future approaches for the diagnosis, risk stratification and therapeutic interventions for pediatric AML.

## Methods

Complete methodological details are provided in the Supplementary Methods. All specimens and clinical data were obtained from patients enrolled on biology studies and clinical trials managed through the Children’s Oncology Group (COG protocols AAML0531 and AAML03P1). Patient samples were sequentially identified, and selected for comprehensive genomic profiling, if adequate amounts of high-quality nucleic acids was available. Patient samples were collected as matched trios: bone marrow aspirates pre-and post-treatment, and matched marrow fibroblasts. Details of sample preparation protocols and clinical annotations and all primary data are available through the TARGET Data Matrix (https://ocg.cancer.gov/programs/target/data-matrix). Whole-genome paired-end sequencing libraries were prepared using the genomic 350-450bp insert Illumina library construction protocol with Biomek FX robot (Beckman-Coulter, USA), sequenced with an average coverage of 30-fold using Illumina HiSeq2500. Sequence files were mapped to the GRCh37 (hg19) genome, and processed to identify single nucleotide variants (SNVs), insertions/deletions (indels), gene fusions, and structural variants. For mRNA-sequencing, extracted RNAs was used to generate cDNAs using the SMART cDNA synthesis protocol with the SMARTScribe reverse transcriptase (Clontech) and resultant libraries were sequenced with 75 bp paired reads using Illumina HiSeq2500. RNA-seq reads were aligned with STAR (version 2.4.2a), and genes annotated in Gencode v18 were quantified with featureCounts (v1.4.3-p1). Fusion genes were detected using FusionCatcher and STAR-Fusion. Resultant variant call files (VCFs) were subsequently aggregated using an integrated script, available from https://github.com/kentsisresearchgroup/TargetInductionFailure. VCFs were parsed to assemble single nucleotide variants, indels, copy number variation, structural variants, and gene fusions in a master table, and filtered to identify high-confidence calls. Normalization and differential expression was done with the Bioconductor package DESeq2. Gene set enrichment analysis was performed using GSEA v2.2.1 plus MSigDB v6.0. All raw sequencing data are available via dbGaP accession numbers phs000465, phs000178 and phs000218, with the processed mutational and expression data published via Zenodo (http://doi.org/10.5281/zenodo.1403737).

## Results

### Genomic landscape of pediatric induction failure AML

A total of 28 patients with primary chemotherapy resistance and failure of induction chemotherapy were studied. The patients were uniformly treated as part of the COG AAML0531 study, having received cytarabine, daunorubicin and etoposide (ADE10+3) chemotherapy. Demographic features of the study cohort are listed in Supplemental Table 1, and are representative of the entire patient cohort, enrolled as part of the AAML0531 study (12). True failure of induction chemotherapy was defined as morphologic persistence of at least 5% of AML bone marrow blasts 28 or more days after therapy initiation, but prior to the second course of induction chemotherapy (12). We used genomic DNA from cultured fibroblasts isolated from the bone marrow as non-tumor germline DNA to identify somatically acquired mutations. Using supervised analysis based on genes currently known to have cancer predisposition potential, we did not identify any apparent pathogenic germline variants in this cohort (Supplemental Table 2). For whole-genome DNA sequencing, we obtained mean coverage of 39 (range 23-69). mRNA and miRNA sequencing data had on average 59% and 19% mapping coverage, respectively.

In agreement with prior studies (11), we found that this cohort of pediatric AML with induction failure had fewer of the mutations commonly observed in adult AML, including *DNMT3A*, *TET2*, *IDH1/2* and others (13). The most commonly called alterations observed in our cohort were rearrangements of *NUP98*, and variants in *WT1*, *RUNX1*, *MLLT10*, *SPECC1*, and *KMT2C*, predominantly as a result of genomic rearrangements and somatic structural variants (Figure 1A). In particular, we identified *NUP98-NSD1* fusions, as well as a number of additional genomic rearrangements, leading to the production of chimeric fusion genes, as evidenced by the combined genomic rearrangements in DNA, and the presence of mRNA sequencing reads in RNA-seq data (Figure 1B). While mutations of *FLT3* and *KMT2C*, and t(8;21), inv(16) and trisomy 8 alterations were the five most common events in the analysis of an unselected cohort of pediatric AML patients (11), these abnormalities were substantially depleted in our induction failure cohort. The relative and unselected enrichment for *NUP98-NSD1* rearrangements and *WT1* mutations in this induction failure cohort is consistent with the reported poor prognosis of these alterations, with a reported 4-year event-free survival of less than 10% (14).

**Figure 1:**
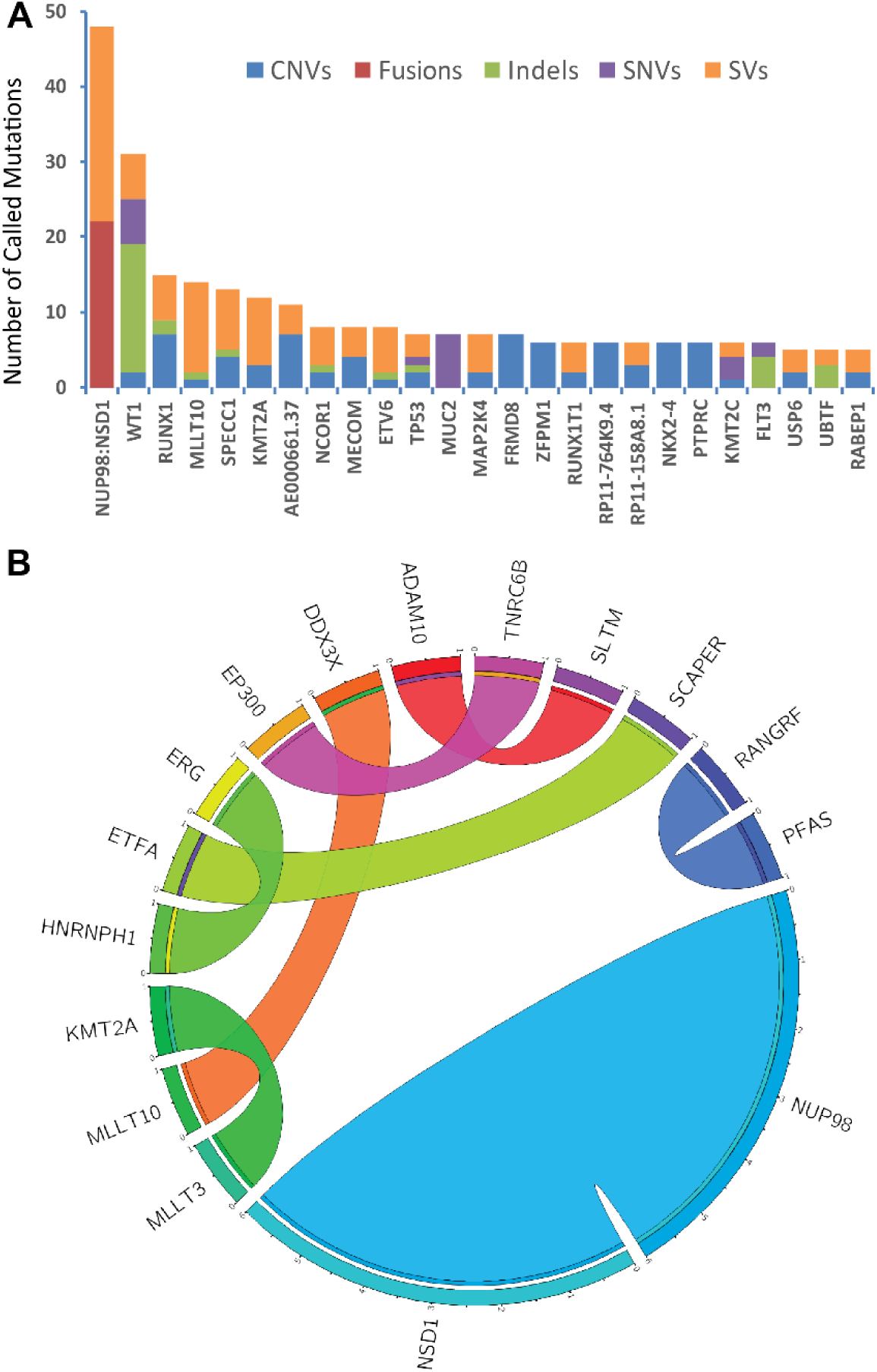
Recurrently mutated genes in pediatric induction failure AML identified by whole-genome and RNA sequencing analysis. **(a)** Top 20 most commonly called mutated genes, with the mutation type indicated by color, observed at diagnosis, enumerated by the number of total calls, independent of patient assignment. **(b)** Circos plot of high-confidence gene fusions, identified from combined analysis of RNA and whole-genome sequencing data, observed at diagnosis. All variants are tabulated as fusions for calls from RNA-seq, structural variants when called from whole-genome sequencing, and both when both the fusion and supporting genomic structural variants match.

### Three genetic subtypes of pediatric AML with primary chemotherapy resistance

Although diverse mutations were observed in our cohort, unsupervised hierarchical clustering was unable to segregate the observed cohort into distinct classes. Therefore, we divided the patients into three groups based on the most common recurrent mutations (Figure 2A). Group 1 (6 patients) was defined by the presence of *NUP98* rearrangements, and additional mutations including *WT1*, *ELF1*, and *FRMD8*. No specimens exhibited chromosomal monosomies or complex karyotypes. Although *FLT3* mutations are often observed in *NUP98-NSD1* leukemias, group 1 did not appear to have *FLT3* mutations. The association of *NUP98* rearrangements with *WT1* mutations in AML induction failure may be due to the functional interaction between these two factors, since patients with both alterations are known to have a much worse prognosis than either alone (11, 14, 15). In addition, we observed an association of *NUP98* rearrangements with deletions of the ETS transcription factor *ELF1* and copy number gain of the gene encoding cell adhesion signaling factor *FRMD8*, both of which have also been observed in myeloid malignancies (16, 17). This association may involve similar cooperating interactions that presumably cause intrinsic chemotherapy resistance. We also identified gain of *MYC* in two patients from group 1, in agreement with prior finding of activating *MYC* mutations in association with *NUP98-NSD1* AML (18).

**Figure 2:**
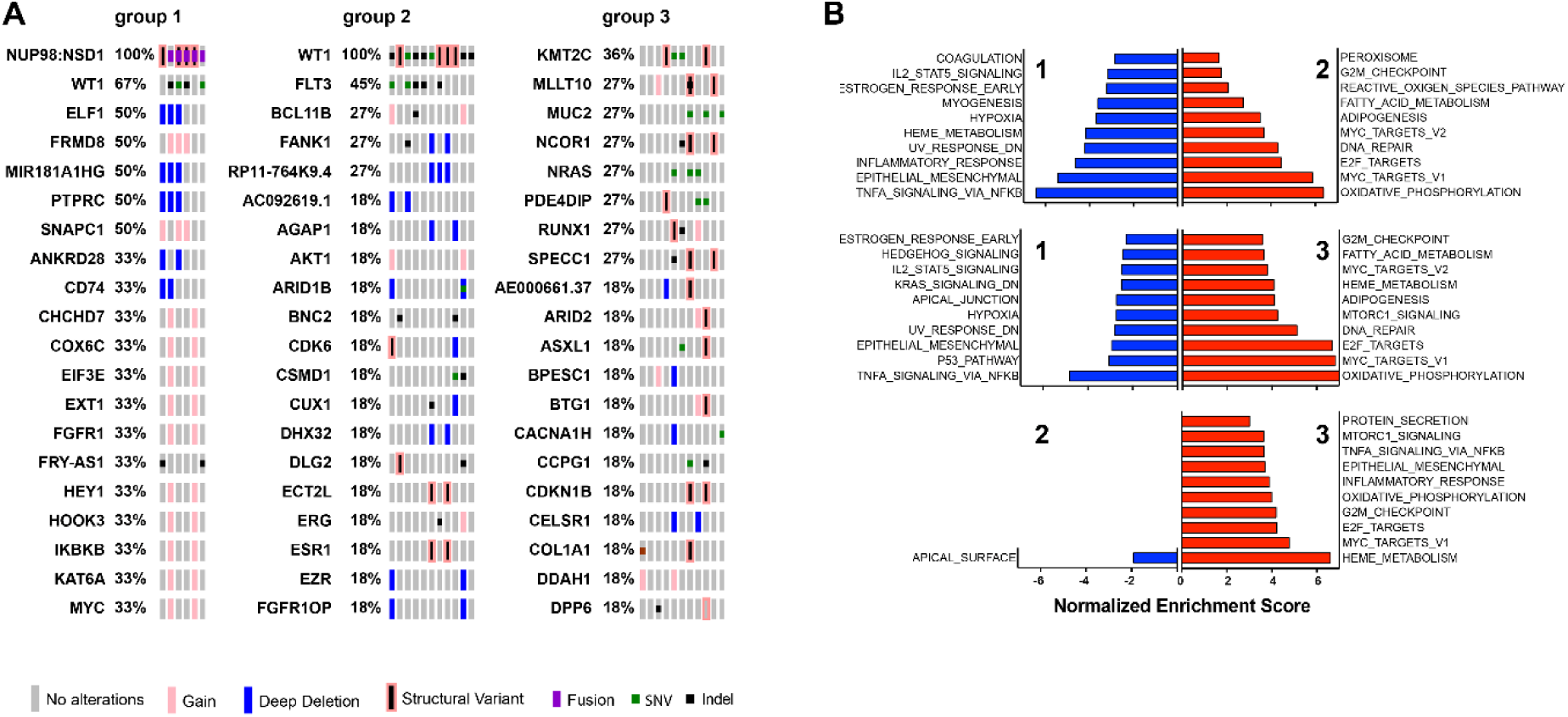
Three groups of pediatric induction failure AML identified by whole-genome and RNA sequencing analysis. **(a)** Tile plot of recurrently mutated genes and gene expression profiles by patient, showing three disease groups, as labeled, with each row listing the mutant gene, and each column representing an individual patient specimen: Group 1, defined by *NUP98* alterations (patients with *NUP98-NSD1* fusions except patient 1 with *NUP98* gain); Group 2, defined by *WT1* mutations, and Group 3, defined by the apparent absence of *NUP98* or *WT1* mutations. **(b)** Gene set enrichment analysis (GSEA) of the three patient groups, listing significantly enriched (red) and downregulated (blue) gene sets, as a function of their normalized enrichment.

Group 2 (11 patients) was defined by the presence of *WT1* mutations without apparent *NUP98* rearrangements, and also involves additional mutations including tyrosine kinase domain (TKD, SNV) and internal tandem duplication (ITD, indel) in *FLT3,* and various other copy number changes and genomic rearrangements. We observed both missense and nonsense *WT1* mutations, consistent with previous reports in AML (19, 20, 21). In addition, group 2 included cases with copy number alterations involving the *BCL11B*, *AKT1*, and *ARID1B* loci, among others (Figure 2A), as well as one patient specimen PATISD with monosomy 7 (Supplemental Table 1). *BCL11B* is a known tumor suppressor gene mutated in refractory forms of T-cell acute lymphoblastic leukemias (T-ALL) (19), including a subtype that may share common origins with refractory AML (20). In addition, both BCL11B and ARID1B are components of the SWI/SNF/BAF chromatin remodeling complex that is disrupted in diverse human cancers (22).

Group 3 (11 patients) was defined by the apparent absence of *NUP98* rearrangements and *WT1* mutations, and instead includes leukemias with mutations of *KMT2C* and *MLLT10* (Figure 2A). *KMT2C* is the tumor suppressor gene that encodes the MLL3 chromatin remodeling factor, that is also inactivated in myeloid malignancies as a result of losses of chromosome 7q (23). Similarly, *MLLT10* is frequently rearranged as part of *KMT2A*/*MLL1* and other chromosomal translocations in acute leukemias, including refractory forms of T-ALL in particular (24) (25). Given the involvement of additional genes and loci recurrently mutated or rearranged in this cohort of patients, it is probable that additional subtypes of chemotherapy resistant disease exist.

Intriguingly, while only one patient in group 1 remained alive at 6 years after therapy, and three patients remained alive in group 2, five survivors were observed in group 3. Though the size of this cohort is not powered sufficiently to detect statistically significant differences in survival (log-rank *p* = 0.39, 0.70, and 0.55 for group 1 vs 2, 1 vs 3, and 2 vs 3 respectively), these results suggest that the apparent diversity of genetic subtypes of induction failure may also be associated with variable clinical outcomes.

Using mRNA sequencing, we analyzed gene expression programs associated with the primary chemotherapy resistant AML, as assessed using gene set enrichment analysis in diagnostic samples (Figure 2B). Unsupervised hierarchical clustering of gene expression profiles did not segregate with the genetically defined groups (Supplemental Figure 1). This suggests that diverse genetic subtypes of induction failure AML may engage common gene expression programs. This notion is consistent with the recent study implicating epigenetic signaling by the transcription factor MEF2C in AML chemotherapy resistance (9).

Lastly, we surveyed microRNA expression in diagnostic samples from this cohort, with the most highly expressed miRNAs listed in Supplemental Table 3. We observed that miR-21 was highly expressed among all 3 subgroups of induction failure patients, consistent with its reported association with inferior clinical outcomes (26). Similarly, we observed high levels of expression of miR-10a, particularly in group 1. miR-10a upregulation has been reported in *NPM1*-mutant AML with associated MDM4 downregulation, potentially interfering with TP53 signaling (27). We also found high expression of miR-103 in group 1 patients, which has been reported to downregulate *RAD51*, leading to dysregulated DNA damage response (28). In addition, we found upregulation of miR-181a in groups 2 and 3, which has been reported to be overexpressed and mediate ATM downregulation in AML cell lines (29). In all, these findings are consistent with the proposed mechanisms of regulation of chemotherapy response by miRNAs in AML (30).

### Diverse models of clonal evolution by induction chemotherapy

We reasoned that exposure to chemotherapy would lead to selection of genetic clones with mutations conferring chemotherapy resistance, and contraction of clones that are susceptible to ADE chemotherapy. Thus, we compared the prevalence of mutations among different patients in specimens collected before and after induction chemotherapy (Figure 3). We found numerous genomic rearrangements and mutations that were significantly increased in prevalence upon induction chemotherapy exposure. For example, we observed that gains of the *FRMD8* locus were present in 7 of 28 (25%) patients at diagnosis, as compared to 20 of 28 (71%) patients post-chemotherapy (two-tailed Fisher’s exact test *p* = 1.1e-3). This suggests that genomic rearrangement involving *FRMD8* or linked genes may contribute to chemotherapy resistance. *FRMD8* encodes a plasma membrane-associated FERM domain that can contribute to Wnt signaling and processing of transmembrane precursors of inflammatory cytokines (31, 32). In addition, increased *FRMD8* gene expression was found to be a marker of poor prognosis in adult AML (33).

**Figure 3:**
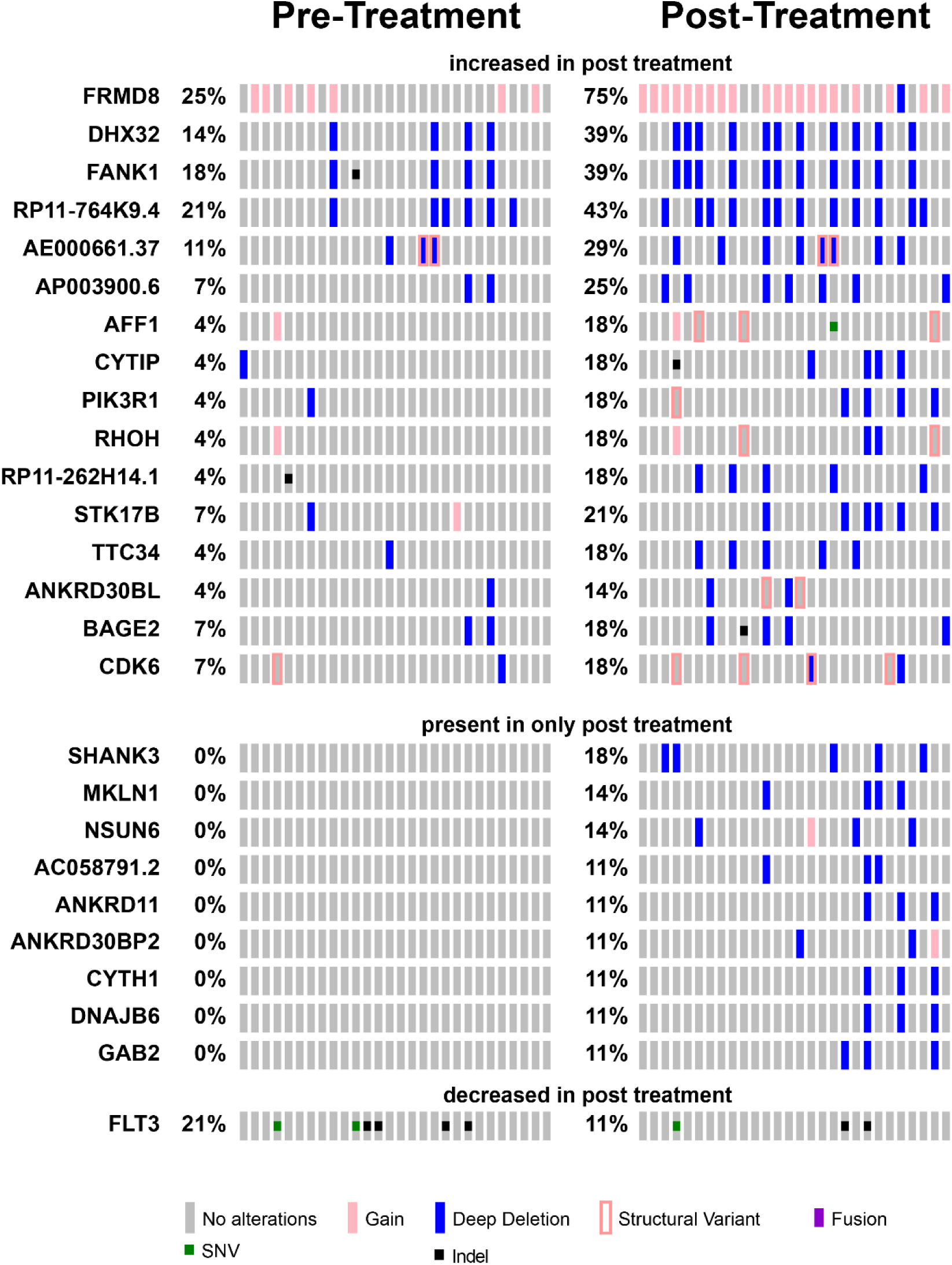
Clonal selection upon chemotherapy treatment. Tile plot showing recurrently mutated genes with changes in apparent allele frequencies upon induction chemotherapy. For *FLT3* mutations, TKD mutations are listed as SNVs, and ITDs as indel mutations.

Mutations and rearrangements of various additional genes with functions in cell adhesion and signaling, including *FANK1*, *PIK3R1*, *SHANK3*, and *MKLN1*, also appear to be selected upon chemotherapy exposure, suggesting that they may also contribute to therapy resistance. In contrast, mutations of *FLT3* exhibited significant depletion upon chemotherapy administration (Figure 3A). Aberrant activation of FLT3 kinase signaling is a known oncogenic event in AML pathogenesis, contributing to the enhanced proliferation and survival of AML cells, and is associated with inferior prognosis when present at sufficiently high allelic frequencies (34-38). Its relative depletion by chemotherapy in AML induction failure suggests that its subclonal evolution in and of itself does not cause chemotherapy resistance. Rather, its activation in combination with specific other pathogenic events as part of distinct clones, such as those with mutations of *WT1* or *NUP98* rearrangements or others (6, 38), may cause resistance to chemotherapy.

In addition to the marked changes in overall clonal architecture associated with induction chemotherapy, we also observed multiple modes of clonal evolution within individual leukemias. In general, induction chemotherapy induced a relative contraction of the AML cell population, as evidenced by the reduction of the apparent variant allele frequencies (VAF) of mutant genes (mean 0.40 versus 0.28 for pre- and post-chemotherapy, respectively, Bonferroni adjusted t-test *p* = 2.6e-4). While VAF contraction was common to most mutations, closer examination of individual patients revealed distinct potential modes of clonal evolution (Figure 4). For example, specimen PASFHK exhibited significant expansion of the *WT1;PTCH1;ZNF785*-mutant clone, and elimination of the *FLT3;SERPIN2*-mutant subclones, upon chemotherapy exposure (Figure 4A). This is consistent with the prior reports of elimination of FLT3-mutant subclones upon AML relapse (5), supporting the proposal that activated FLT3 contributes to chemotherapy resistance only when present with specific cooperating mutations, such as *WT1*. For specimen PATJMY, we observed evolution of a new loss-of-function nonsense mutation of *CHMP6*, which emerged either upon chemotherapy exposure or was selected as a pre-existing subclone, present at less than 2% fraction at diagnosis, given the 50-fold sequencing coverage for *CHMP6* (Figure 4B). Reduced *CHMP6* gene expression has been associated with inferior survival of elderly AML patients (39), and its function in endosomal cell surface receptor recycling may contribute to chemotherapy resistance (40). In agreement with prior reports (7), specimen PASTZK exhibited subclonal evolution of mutant *TP53* at diagnosis, which led to its clonal expansion in combination with clonal *PHF6* mutation upon chemotherapy administration, in contrast to mutation of *NRAS* which remained subclonal (Figure 4C). Finally, specimen PARXYR exhibited relative contraction of the *WT1;PTPN11*-mutant subclone, and relative expansion of the *GPR137B*-mutant subclone that additionally acquired a *CD82* mutation (Figure 4D). Other leukemias showed similar subclonal composition pre- and post-treatment, such as for specimen PARBTV, which demonstrated the likely pathogenic *IDH2* R172K (VAF 0.58 pre and 0.45 post) and *H3F3A* K27M mutations (0.46 pre and 0.54 post). These findings demonstrate distinct modes of clonal selection upon chemotherapy exposure, which are expected to inform future targeting of specific molecular mechanisms to overcome or block chemotherapy resistance.

**Figure 4:**
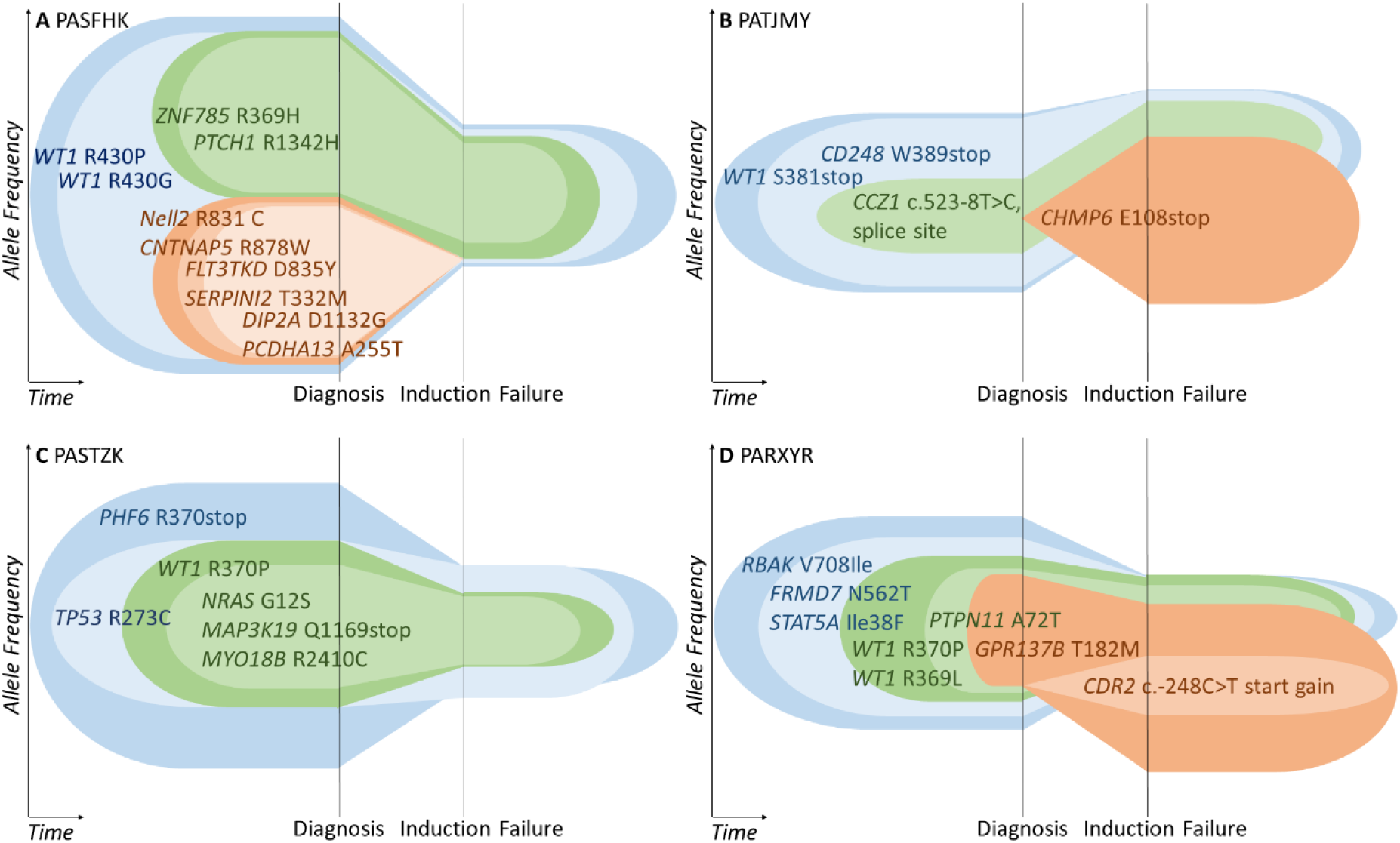
Mutant allele frequencies suggest diverse modes of clonal evolution upon chemotherapy exposure. Variant allele frequencies (VAF) in four specimens with *WT1* mutant clones are represented as the height of color-coded clones, with full height of allele frequency axis corresponding to a mutant allele frequency of 1. Simplest models of clonal architecture are shown, with additional models possible. Each hypothesized subclone is represented by a different color. **(a)** Specimen PASFHK in Group 2, largest VAF at diagnosis was 0.96 for *WT1* R430P; largest VAF at induction failure was 0.38 for *WT1* R430P. **(b)** Specimen PATJMY in Group 1, largest VAF at diagnosis was 0.53 for *WT1* S381, largest VAF at induction failure was 0.47 for *CHMP6* E108stop. **(c)** Specimen PASTZK in Group 2, largest VAF at diagnosis was 0.77 for *PHF6* R370stop, largest VAF at induction failure was 0.37 for *TP53* R273C. **(d)** Specimen PARXYR in Group 1, largest VAF at diagnosis was 0.57 for *RBAK* V708I, largest VAF at induction failure was 0.43 for *GPR137B* T182M.

## Discussion

Our study defines the genomic landscape of pediatric AML with primary chemotherapy resistance and failure of induction remission therapy. Importantly, primary chemotherapy resistant pediatric AML involves multiple distinct genetic mechanisms. Most notably, we found substantial prevalence of structural rearrangements, at least some of which are associated with the expression of chimeric fusion genes. In particular, we observed at least three distinct genetic groups of patients with induction failure, including those with *NUP98* rearrangements, somatic mutations of *WT1*, *ELF1*, *KMT2C*, *MLLT10,* and additional recurrent gene mutations, fusions, and structural rearrangements, some of which have been observed in other malignancies. Given the known technical challenges with the detection of genomic rearrangements and gene fusions (41), it is possible that additional pathogenic structural variants or chimeric gene fusions may contribute to AML and primary chemotherapy resistance.

In our prior study of primary chemotherapy resistance, we identified individual mutations of *ASXL1, SETBP1* and *RELN* to be significantly enriched in a subset of pediatric AML with primary induction failure (8). Insofar as *ASXL1* and *WT1* mutations are mutually exclusive in both pediatric and adult AML, the prevalence of *WT1* mutations and absence of apparent *ASXL1* mutations in our current cohort suggests that additional genetic mechanisms of primary chemotherapy resistance likely exist. Our results also suggest that varied genetic mechanisms of chemotherapy resistance may converge on coherent gene expression programs, at least insofar as they cannot be statistically decomposed using matrix factorization used as part of this gene set enrichment analysis.

Importantly, our study identified distinct combinations of mutations that appear to be associated with primary chemotherapy resistance. In particular, we observed an association between *NUP98-NSD1* fusions and mutations of *WT1*, *ELF1* and *FRMD8*, suggesting possible cooperativity in their pathogenic functions. Similarly, we observed an association between *WT1* mutations and rearrangements of *BCL11B* and *ARID1B* loci, both of which encode components of the SWI/SNF/BAF chromatin remodeling complex. Notably, *BCL11B* is recurrently mutated in refractory forms of T-ALL, which may share common origins with subsets of AML (20). Evidently, these combinatorial mechanisms in pediatric AML are distinguished from other mechanisms of chemotherapy resistance, such as inactivation of *TP53* in adult AML (7).

Our study identified additional mutations associated with pediatric primary chemotherapy resistance. This includes loss-of-function mutations of *KMT2C*, which encodes a component of the MLL3 chromatin remodeling complex, potentially similar to the deletions of chromosome 7q observed in high-risk AML that involve this locus and have been found to confer susceptibility to epigenetic therapies (23). We also observed deletions of *MLLT10*, which is recurrently rearranged as gene fusions in subsets of T-ALL. Insofar as MLLT10 is a cofactor of the DOT1L methyltransferase, this may be associated with the susceptibility to emerging DOT1L methyltransferase inhibitors such as pinometostat (EPZ-5686), which will need to be tested in future studies. Additional mutations associated with primary chemotherapy resistance may be found in larger studies. For instance, the presence of likely pathogenic *IDH2* R172K and *H3K27M* K27M mutations in one specimen in our cohort suggests additional potential mechanisms of chemotherapy resistance (42-45), which may confer susceptibility to emerging therapies such as the IDH inhibitor enasidenib (AG-221) for example.

Our findings also suggest that the diversity of genetic chemotherapy resistance mechanisms may be associated with variable outcomes of intense combination chemotherapy in AML. Importantly, increase in the apparent prevalence and allelic frequency of genetic clones with mutations of *FRMD8, FANK1, PIK3R1, WT1* and others indicate that these alleles, in cooperation with *NUP98-NSD1* and other initiating mutations, may directly cause chemotherapy resistance. In contrast, subclonal mutations of *FLT3, PTPN11* and *NRAS* were reduced or eliminated by chemotherapy, suggesting that these secondary mutations in and of themselves do not cause chemoresistance. Indeed, subclonal mutations of *FLT3* or *NRAS* were not significantly associated with primary chemotherapy resistance in our prior study (8). Diverse genetic mechanisms of chemotherapy resistance may be associated with clonal evolution (5, 6), as also evidenced by our findings (Figures 3 and 4). On the other hand, common gene expression programs may be associated with shared molecular dependencies, substantiating the development of targeted therapies, as recently evidenced by molecular therapy of MEF2C in chemotherapy resistant AML (46).

In all, our study demonstrates that primary chemotherapy resistance and failure of induction chemotherapy in pediatric AML is associated with multiple genetic mechanisms, and exhibits diverse clonal dynamics, dependent on distinct combinations of mutations. Future functional studies will be needed to assess the mechanisms of cooperativity among the observed chemotherapy-associated mutations and their specific pharmacologic targeting. Similarly, additional studies will be needed to define the prognostic significance of the observed chemotherapy-associated mutations. This is expected to delineate molecular mechanisms of primary chemotherapy resistance in pediatric AML, which should inform improved approaches for its diagnosis and therapy.

## Supporting information

## Acknowledgements

This paper is dedicated to the memory of Dr. Robert Arceci. This work was supported by the NCI U10 CA98543 (TARGET), U24 CA114766 (COG), U10 CA180886, U10 CA098413, U10 CA180899, P30 CA008748, R01 CA204396, by the Damon Runyon-Richard Lumsden Foundation Clinical Investigator and St. Baldrick’s Foundation Arceci Innovation Awards (A.K.), and the Charles E. Trobman Scholarship (N.M.). We thank T. Davidsen and P. Gesuwan for their support of the TARGET Data Coordinating Center, and Alejandro Gutierrez, Gila Spitzer, and Maria Luisa Sulis for comments on the manuscript.

## Notes

**Conflicts of Interest.** A.K. is a consultant for Novartis.

