## Supplementary material for "Genetic mechanisms of primary chemotherapy resistance in pediatric acute myeloid leukemia: A report from the TARGET initiative"

### Supplemental Figures and Tables

#### Supplemental Table 1. Description of patient cohort.

| TARGET USI | Gender | Race | Age (days) at diagnosis | EFS (days) | OS (days) | Vital Status | Year Diagnosed | Year of last follow-up | Assigned Group | Cytogenetics |
|---|---|---|---|---|---|---|---|---|---|---|
| PAMYMA | Male | White | 450 | 28 | 378 | Dead | 2004 | 2005 | 1 | 46,Y,t(X;11)(q13;p15.1)[18]/46,XY[2] |
| PANZLR | Female | Black | 4793 | 28 | 74 | Dead | 2005 | 2005 | 1 | 47,XX,+8[20] |
| PARBTV | Male | White | 6273 | 77 | 444 | Dead | 2007 | 2008 | 3 | 46,XY[20] |
| PARHRS | Male | White | 3636 | 82 | 2616 | Alive | 2007 | 2014 | 2 | 47,XY,+4,t(15;21)(q22;q22)[11]/46,XY[10] .ish AML1sp |
| PARLSL | Male | White | 4456 | 1077 | 1077 | Alive | 2007 | 2010 | 3 | 46,XY[20] |
| PARNAW | Female | White | 430 | 66 | 457 | Dead | 2007 | 2009 | 2 | 46,XX[28] |
| PARXYR | Female | White | 5346 | 492 | 2200 | Alive | 2008 | 2014 | 1 | 46,XX[20] |
| PARZIA | Female | Asian | 6089 | 77 | 77 | Dead | 2008 | 2008 | 3 | 47,XX,+10[7]/46,XX[13] |
| PASDKZ | Female | Asian | 1875 | 188 | 188 | Dead | 2008 | 2009 | 1 | 46,Y,t(X;10)(p11.2;p11.2),add(17)(p11.2)[13]/46,XY[6] |
| PASDXR | Male | White | 3440 | 33 | 276 | Dead | 2008 | 2009 | 1 | 47,XY,+8[20] |
| PASFHK | Male | White | 2164 | 85 | 1655 | Alive | 2008 | 2013 | 2 | 46,XY,add(14)(q32)[18]/45,idem,psu dic(9;9)(p11;p22)[2] |
| PASFJJ | Female | White | 3836 | 2205 | 2205 | Alive | 2008 | 2014 | 2 | 46,XX,inv(2)(p13q22)mat[25] |

| | | | | | | | | | | |
|---|---|---|---|---|---|---|---|---|---|---|
| PASFLG | Male | White | 3971 | 69 | 694 | Dead | 2008 | 2010 | 2 | Unknown |
| PASIGA | Male | White | 318 | 1947 | 1947 | Alive | 2008 | 2014 | 3 | 46,XY,t(9;11)(p22;q23)[15]/47,idem,+19[2]/46,XY[4] |
| PASLZE | Female | Black | 3711 | 84 | 707 | Dead | 2009 | 2011 | 3 | 46,XX,t(8;21)(q22;q22)[20] |
| PASNKZ | Female | White | 6491 | 195 | 195 | Dead | 2009 | 2009 | 3 | 46,XX,del(7)(q22)[15]/46,XX[5] |
| PASSLT | Male | Unknown | 5405 | 36 | 87 | Dead | 2009 | 2009 | 3 | 46,XY,t(10;11)(p13;q14),der(17)t(17;17)(p13;q21),del(12)(p12),+2mar[4]/48,idem,t(13;15)(q32;q13)[11]/48,idem,add(3)(p12),der(4)inv(4)(q12q25)add(4)(q25),-del(12),der(13)(13pter->13q34::?3p21->3p13::4q25->4qter)[5] |
| PASTZK | Male | Black | 2145 | 58 | 102 | Dead | 2009 | 2009 | 2 | 46,Y,t(X;10)(p11.2;p11.2),add(17)(p11.2)[13]/46,XY[6] |
| PASVJS | Female | White | 1913 | 71 | 1298 | Dead | 2009 | 2013 | 2 | 46,XX,[20] |
| PASYEJ | Female | White | 889 | 69 | 988 | Dead | 2009 | 2012 | 3 | 46,XX[30] |
| PASYWA | Male | White | 5181 | 79 | 295 | Alive | 2009 | 2010 | 3 | 46,XY[20] |
| PATAIJ | Female | Black | 417 | 57 | 1660 | Alive | 2009 | 2014 | 3 | 46,XX,cryp ins(10;11)(p12;q23q23),inv(17)(p13.1q11.2)[20] |
| PATHIU | Male | White | 4859 | 374 | 454 | Dead | 2010 | 2011 | 2 | 47,XY,+8[14]/46,XX[6] |
| PATISD | Male | White | 3901 | 46 | 148 | Dead | 2010 | 2010 | 2 | 45,XY,t(3;3)(q21;q26),-7[20] |
| PATJMY | Male | White | 1799 | 77 | 319 | Dead | 2010 | 2011 | 1 | 46,XY[20] |
| PATKBK | Male | White | 31 | 1684 | 1684 | Alive | 2010 | 2014 | 3 | 46,XY,t(1;22)(p13;q13)[7]/46,XY[13] |
| PATKKJ | Female | Unknown | 8581 | 451 | 451 | Dead | 2010 | 2011 | 2 | 46,XX,add(9)(p13),+21[16]/46,XX[2] |
| PATKWH | Male | White | 6008 | 504 | 701 | Dead | 2010 | 2012 | 2 | 46,XY[20] |

**Supplemental Table 2. Observed germline variants.**

| 1000G_AF | ExAC_AF | ID | CHROM | POS | REF | ALT | Mutation Assessor_pred | SNPEFF_AA_CHANGE | SNPEFF_CDS_CHANGE | SNPEFF_EFFECT | SNPEFF_FUNCTIONAL_CLASS | SNPEFF_GENE_BIOTYPE | SNPEFF_GENE_NAME | SNPEFF_IMPACT | SNPEFF_TRANSCRIPT_ID | Sample | ACMG |
|---|---|---|---|---|---|---|---|---|---|---|---|---|---|---|---|---|---|
| 2.00E-04 | 1.07E-04 | rs116788608 | 7 | 6035211 | T | C | M | p.Asp286Gly | c.857A>G | NON_SYNONYMOUS_CODING | MISSENSE | protein_coding | PMS2 | MODERATE | ENST00000265849 | PAMXZY | VUS |
| 2.40E-03 | 4.55E-04 | rs115574135 | 7 | 116409777 | C | T | N | p.His906Tyr | c.2716C>T | NON_SYNONYMOUS_CODING | MISSENSE | protein_coding | MET | MODERATE | ENST00000318493 | PANZLR | VUS |
| 1.40E-03 | 7.50E-04 | rs148590073 | 11 | 108106435 | A | G | N | p.Ile124Val | c.370A>G | NON_SYNONYMOUS_CODING | MISSENSE | protein_coding | ATM | MODERATE | ENST00000278616 | PANZLR | LB |
| 1.60E-03 | 2.55E-04 | rs35423758 | 16 | 68847433 | C | T | | p.Ala400Val | c.1199C>T | NON_SYNONYMOUS_CODING | MISSENSE | nonsense_mediated_decay | CDH1 | MODERATE | ENST00000566510 | PANZLR | VUS |
| 2.60E-03 | 1.58E-03 | rs150600452 | 13 | 32949533 | A | G | | p.Ile62Val | c.184A>G | NON_SYNONYMOUS_CODING | MISSENSE | nonsense_mediated_decay | BRCA2 | MODERATE | ENST00000528762 | PARHRS | VUS |
| 2.00E-04 | 1.79E-03 | rs77724903;COSM1159820 | 10 | 43613908 | A | T | N | p.Tyr791Phe | c.2372A>T | NON_SYNONYMOUS_CODING | MISSENSE | protein_coding | RET | MODERATE | ENST00000355710 | PARLSL | VUS |
| 9.98E-04 | 1.99E-03 | rs118101777 | 15 | 90630704 | C | T | H | p.Arg261His | c.782G>A | NON_SYNONYMOUS_CODING | MISSENSE | protein_coding | IDH2 | MODERATE | ENST00000330062 | PARLSL | LB |
| 9.98E-04 | 1.61E-03 | rs143638171;COSM30751 | 5 | 112174677 | T | C | N | p.Leu1129Ser | c.3386T>C | NON_SYNONYMOUS_CODING | MISSENSE | protein_coding | APC | MODERATE | ENST00000257430 | PASDXR | LB |
| 2.00E-03 | 2.49E-03 | rs3219496 | 1 | 45795043 | G | T | M,M | p.Leu529Met | c.1585C>A | NON_SYNONYMOUS_CODING | MISSENSE | protein_coding | MUTYH | MODERATE | ENST00000450313 | PASFHK | LB |
| 4.79E-03 | 1.21E-03 | rs386833391;rs567584401;COSM4170075 | 5 | 112174750 | TGAA | T | | p.Glu1157del | c.3468_3470delAGA | CODON_CHANGE_PLUS_CODON_DELETION | | protein_coding | APC | MODERATE | ENST00000257430 | PASLZE | LB |
| 9.98E-04 | 6.42E-04 | rs2227971;rs138482490 | 9 | 98231061 | G | A | M | p.Ala741Val | c.2222C>T | NON_SYNONYMOUS_CODING | MISSENSE | protein_coding | PTCH1 | MODERATE | ENST00000331920 | PASVJS | LB |
| 2.00E-04 | 2.02E-03 | rs1800059 | 11 | 108170506 | A | C | N | p.Ser1691Arg | c.5071A>C | NON_SYNONYMOUS_CODING | MISSENSE | protein_coding | ATM | MODERATE | ENST00000278616 | PATISD | LB |

A

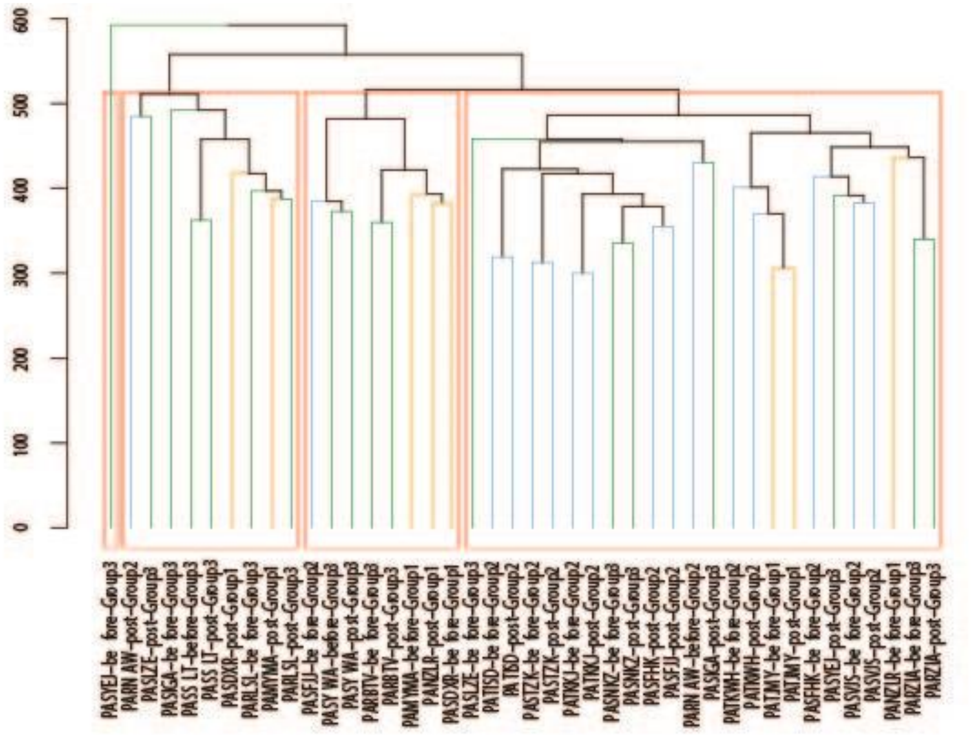

B

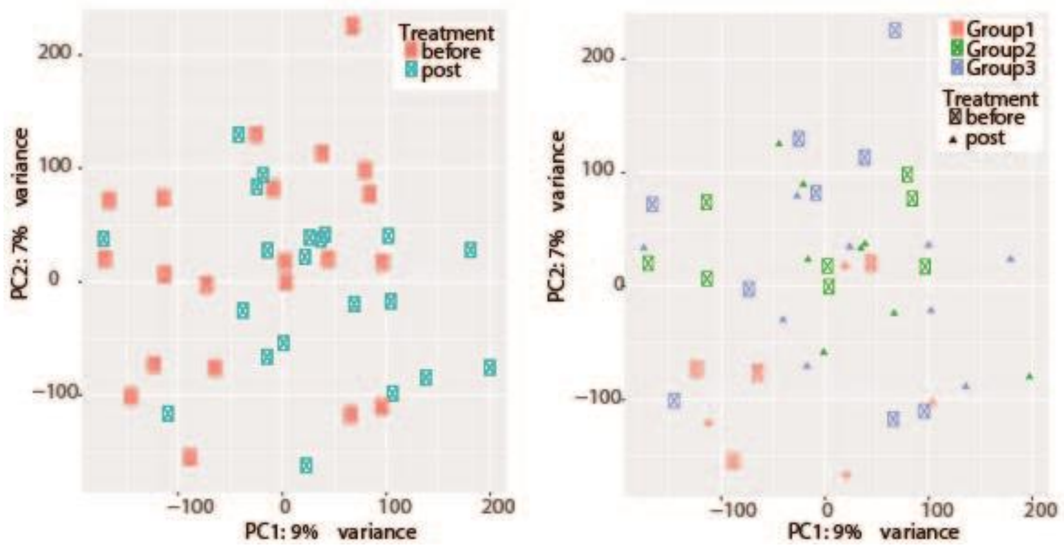


**Supplemental Figure 1. Unsupervised clustering (A) and principal component analysis (B) suggest a lack of clear differentiation of expression in genes among genetically defined groups.** Analysis was performed using log2 of normalized counts obtained from DESeq2 (v1.14.1). Euclidean distances were calculated using base R's dist function, and clustering obtained using base R's hclust function on the Euclidean distances. Base R's cutree function was used to divide the samples into 4 groups based on the clustering results. This resulted in 3 main groups, plus a fourth group with a single sample.

**Supplemental Table 3. Highly expressed miRNA pre-treatment.** Tabulated are the top five most highly expressed miRNA present in > 2 patients in each group pre-treatment. Percentage of how many patients for whom the indicated miRNA was among the top five is given in parentheses.

| Group 1 | Group 2 | Group 3 |
|---|---|---|
| miR-21 (100%) | miR-92a (100%) | miR-92a (100%) |
| miR-103a (100%) | miR-21 (89%) | miR-21 (89%) |
| miR-10a (60%) | miR-101 (78%) | miR-101 (56%) |
| miR-92a (60%) | miR-25 (56%) | miR-25 (44%) |
| miR-101 (60%) | miR-103a (56%) | miR-103a (44%) |
| miR-25 (40%) | miR-181a (56%) | miR-181a (33%) |
| | miR-10a (33%) | miR-10a (22%) |

### Supplemental Methods

**Sample preparation.** Comprehensive details regarding sample preparation is available in the TARGET sample matrix (https://ocg.cancer.gov/programs/target/data-matrix). DNA and RNA were extracted from Ficoll-enriched cryopreserved samples from the COG biorepository using the AllPrep Extraction Kit (Qiagen).

**Marrow Fibroblast Culture.** Bone marrow cells in freezing media were thawed quickly in a 37°C water bath and transferred to a 50ml tube. 1ml of warm Chang media was added to the empty cell vial to wash remaining cells, then transferred to the 50ml tube. Cells rested for 3 minutes then 4 more ml of media were added to the tube. Cells rested another 3 minutes then 8 more ml of media were added. Cells rested another 3 minutes, then were spun for 5 minutes at 1200 rpm. Supernatant was discarded and cells were resuspended in 4 ml of fresh Chang media. Cells were counted and then transferred to a T75 flask. 10 more ml of media were added to flask before being placed in a humidified 37°C, 5% $CO_2$ incubator. Growing fibroblasts were checked 3 days after thawing, old media was removed from flask and discarded while 15ml warm Chang media was added to flask before re-incubation. Cells were checked and media changed every 3-4 days. Cells were split when confluent at 70-80%.

**Whole genome sequencing.** Genomic DNA was fragmented by Covaris E210 sonication and a paired-end sequencing library was prepared following the BC Cancer Agency's Genome Sciences Centre 96-well Genomic ~350bp-450bp insert Illumina Library Construction protocol with Biomek FX robot (Beckman-Coulter, USA). DNA was purified in a 96-well microtitre plate using Ampure XP SPRI beads and was subject to end-repair, and phosphorylation by T4 DNA polymerase, Klenow DNA Polymerase, and T4 polynucleotide kinase respectively in a single reaction, followed by cleanup using Ampure XP SPRI beads and 3' A-tailing by Klenow fragment (3' to 5' exo minus). Picogreen quantification was performed to determine the amount of Illumina PE adapters used for ligation. The adapter-ligated products were purified using Ampure XP SPRI beads, then PCR-amplified with Phusion DNA Polymerase (Thermo Fisher Scientific Inc. USA) using Illumina's PE indexed primer set, with cycle conditions: 98˚C for 30sec followed by 6 cycles of 98˚C for 15 seconds, 62˚C for 30 sec and 72˚C for 30 sec, and a final extension at 72˚C for 5min. The PCR products were purified using Ampure XP SPRI beads, and checked with Caliper LabChip GX for DNA samples using the High Sensitivity Assay (PerkinElmer, Inc. USA). PCR product of desired size range was gel purified (8% PAGE or 1.5% Metaphor agarose in an in-house custom-built robot), and the DNA quality was assessed and quantified using an Agilent DNA 1000 series II assay and Quant-iT dsDNA HS Assay Kit using Qubit fluorometer (Invitrogen), then diluted to 8nM. The final concentration was confirmed by Quant-iT dsDNA HS Assay prior to Illumina Sequencing.
The adapter and sequencing primers used were:

Adapter 5' CAAGCAGAAGACGGCATACGAGAT**NNNNNN**CGGTCTCGGCATTCCTGCTGAACCGCTCTTCCGATCT

Adapter 3' AATGATACGGCGACCACCGAGATCTACACTCTTTCCCTACACGACGCTCTTCCGATCT

| | |
|---|---|
| Seq read 1 primer | ACACTCTTTCCCTACACGACGCTCTTCCGATCT |
| Seq read 2 (index) primer | GATCGGAAGAGCGGTTCAGCAGGAATGCCGAGACCG |
| Seq read 3 primer | CGGTCTCGGCATTCCTGCTGAACCGCTCTTCCGATCT |

NNNNNN = one of 96 fault tolerant indices.

**mRNA sequencing.** For each sample, approximately 10 ng of total RNA was processed using the SMART cDNA synthesis protocol including SMARTScribe Reverse Transcriptase (Clontech, #639536). This method deploys a modified oligo(dT) primer to prime the first strand synthesis reaction and a template switching mechanism to generate full-length single-stranded cDNAs containing the complete 5' end of the mRNA as well as universal priming sequences for end-to-end amplification during 20 cycles of PCR. The amplified cDNA was subject to Illumina paired-end library construction using NEBNext paired-end DNA sample Prep Kit (NEB, E6000B-25). Libraries were sequenced with paired 75 bp reads on Illumina HiSeq2500 instruments.

*WGS QC Metrics*: We ran Fastqc pre-alignment and post-alignment QC with Picard/GATK/samtools, looking at coverage, duplicate rates, insert size distributions, and base call quality (per read and per base, both pre- and post-recalibration).

**Data preprocessing.** Before calling, tumor and matched normal DNA sequencing data were preprocessed using the Broad "best practices" pipeline, which includes aligning reads to the GRCh37 human reference genome using the Burrows-Wheeler Aligner (BWA) (Li and Durbin, 2009), marking of duplicate reads by the use of Picard tools (http://picard.sourceforge.net); realignment around indels (done jointly for all samples derived from one individual, e.g. tumor and matched normal samples, or normal, primary and metastatic tumor trios) and base recalibration via Genome Analysis Toolkit (GATK) (McKenna *et al.*, 2010).

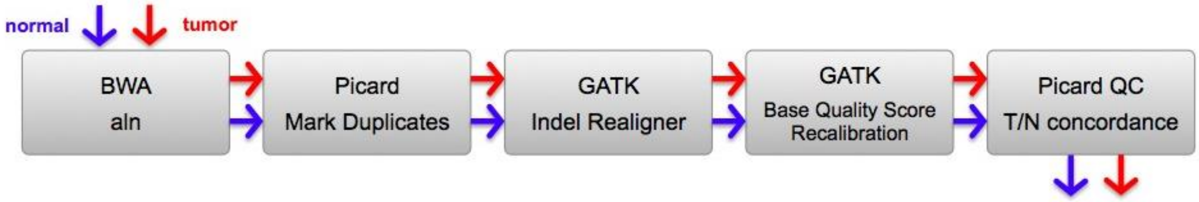


**Methods Figure 1. Pre-processing pipeline.**

**Calling SNVs and indels.** We used the union of somatic SNVs called by muTect (Cibulskis *et al.*, 2013), Strelka (Saunders *et al.*, 2012) and LoFreq (Wilm *et al.*, 2012) and the union of indels called by Strelka, and somatic versions of Pindel (Ye *et al.*, 2009) and Scalpel (Narzisi *et al.*, 2014).

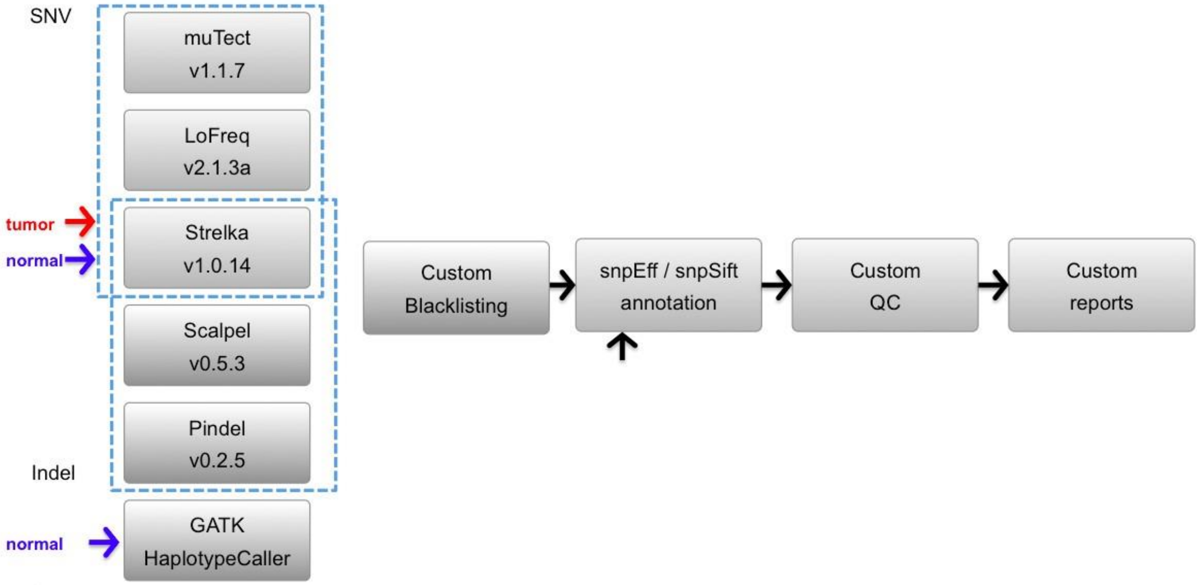


**Methods Figure 2**: **NYGC somatic SNV/indel pipeline.**

The choice of SNV callers was based on internal benchmarking of individual and combinations of callers on a synthetic virtual tumor created by spiking reads from two HapMap samples in a way that mimics somatic variants with predefined variant allele frequencies (Cibulskis *et al.*, 2013). The choice of indel callers was based on internal benchmarking on synthetic data from the DREAM challenge (Ewing *et al.*, 2015).

We also identified germline calls in a panel of cancer risk genes (APC, ATM, BARD1, BMPR1A, BRCA1/2, BRIP1, CDH1, CDK4, CDKN2A, CHEK2, CYLD, EPCAM, IDH1/2, MEN1, MET, MLH1, MSH2/6, MUTYH, NBN, NF1/2, PALB2, PMS1/2, PRKAR1A, PTCH1, PTEN, RAD51C/D, RB1, RET, SDHAF2, SDHB/C/D, SMAD4, STK11, TP53, TSC1/2, VHL, WRN, WT1), made by the use of GATK HaplotypeCaller. Germline variants were filtered for population frequency < 0.005, with low and modifier variants removed, pseudogenes removed, and silent coding variants removed.

**Calling CNVs and SVs.** Structural variants (SVs), such as deletions and amplifications as well as copy-neutral genomic rearrangements were detected by the use of multiple tools (NBIC-seq, Crest, Delly, BreakDancer) that employ complementary detection strategies, such as inspecting read depth within genomic windows, analyzing discordant read pairs, and identifying breakpoint-spanning split reads.

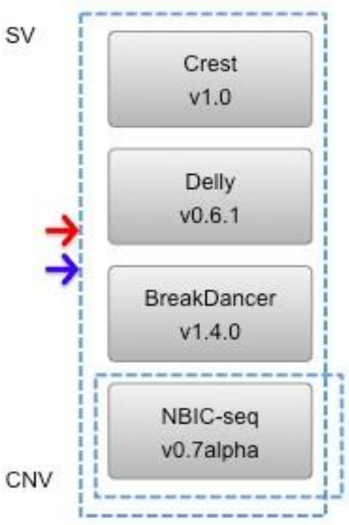


**Methods Figure 3: Somatic CNV/SV pipeline.**

**Filtering SNVs and indels.** We use a multi-step filtering process summarized in Methods Figure 4.

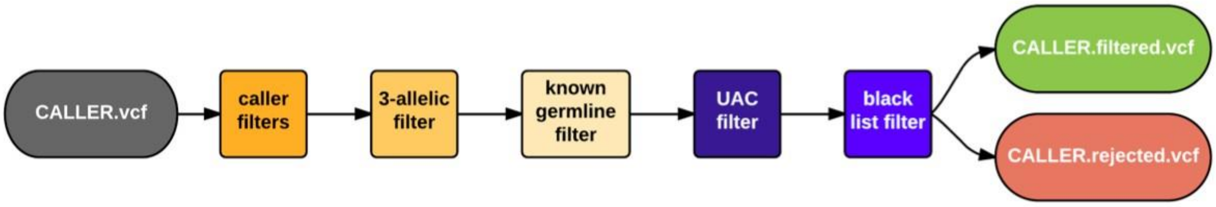


**Methods Figure 4: Custom multi-step SNV/indel filtering**

**Default caller filters.** SNVs and indels were filtered using the default filtering criteria as natively implemented in each of the callers. For Pindel and Scalpel (native germline callers) we used custom in-house scripts for filtering. For each caller we keep these variants:

- LoFreq: FILTER=PASS
- muTect: variants with “PASS” in the filter field of the VCF file, which is equivalent to “KEEP” in the text file
- Strelka: FILTER=PASS
- Pindel: FILTER=PASS
- Scalpel: FILTER=PASS

**Common germline variants.** The resulting set of SNVs and indels was further filtered with common variants seen at MAF ≥ 5% in DNMT3A, TET2, JAK2, ASXL1, TP53, GNAS, PPM1D, BCORL1 and SF3B1 genes (see Xie *et al.*, 2014) and with MAF ≥ 1% elsewhere in the genome, as reported in the 1000 Genomes Project release 3 (1000 Genomes Project Consortium, 2012) and the Exome Aggregation Consortium (ExAC) server (http://exac.broadinstitute.org), because these are very unlikely to be important in cancer.

**UAC filter.** Because callers often return different ref/alt allele counts for the same variant we introduced unified allele counts (UAC). Computation of UAC is based on the bam-readcount tool (Larson *et al.*, 2012). For each variant we generate 4 values that are independent of callers: tumor-ref, tumor-alt, normal-ref, normal-alt. If the tumor_VAF < normal_VAF we discarded the variant.

**Artifacts.** We removed a subset of artifactual calls by the use of a blacklist created by calling somatic variants on 16 random pairings of 80x/40x in-house sequenced HapMap WGS data.

**Annotation and prioritization of SNVs and indels.** Variants were annotated for their effect (non-synonymous coding, nonsense, etc.) using snpEff (Cingolani *et al.*, 2012) based on human genome annotations from ENSEMBL. We further annotate the variants via snpEff, snpSift and GATK VariantAnnotator module with information from COSMIC (Forbes *et al.*, 2012), 1000 Genomes Project, ExAC, CIViC (Clinical Interpretation of Variants in Cancer, https://civic.genome.wustl.edu), and UniProt (http://www.uniprot.org). We returned variant prioritization scores for coding changes based on CHASM (Carter *et al.,* 2009), MutationAssessor (Reva *et al.*, 2011) and FATHMM Somatic (Shihab *et al.,* 2013).

**SV merging.** We merged and annotated SVs called by Crest, Delly and BreakDancer using the BEDPE format. Two SV calls were merged if they shared at least 50% reciprocal overlap (for intra-chromosomal SVs only), their predicted breakpoints were within 300bp of each other and breakpoint strand orientations match for both breakpoints. Thus, merging was done independent of which SV type was assigned by the SV caller (a classification that we found to be unreliable and variable from caller to caller).

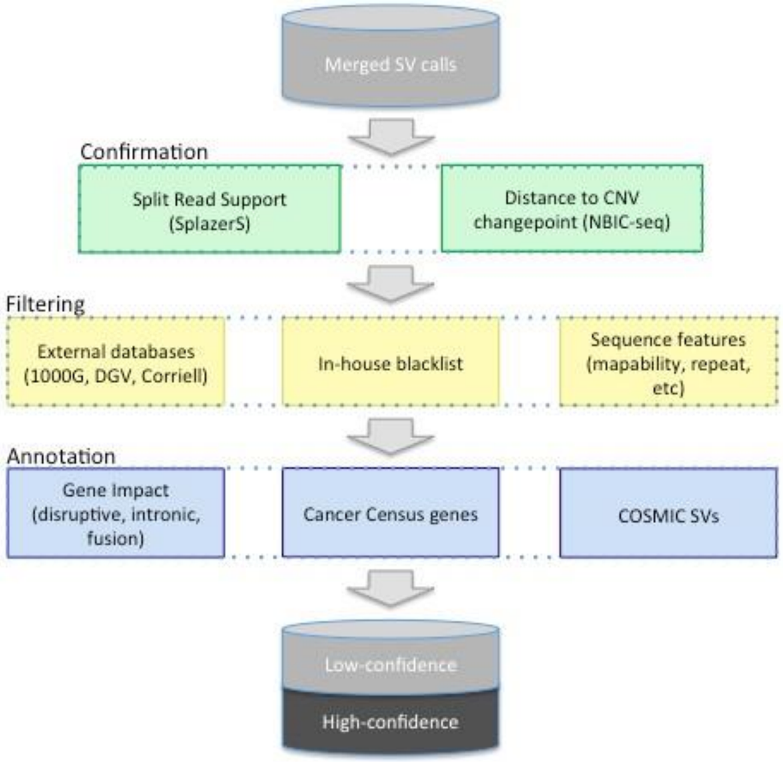


**Methods Figure 5: Somatic CNV/SV filtering and annotation pipeline.**

**Additional SV confirmation.** After merging, we annotated each SV with the closest CNV changepoint as detected by NBIC-seq from read depth signals. This added confidence to true SV breakpoints that were not copy-neutral. Additionally, we used an independent sensitive split read check for each breakpoint using SplazerS. Apart from adding confidence and basepair precision to the breakpoint, this step also helped to remove remaining germline SVs also found in the normal.

**SV filtering.** Some SV callers still suffer from large numbers of false positives; those are often due to germline SVs overlooked in the normal, e.g. because of low coverage or an unmatched normal, or systematic artifacts due to mapping ambiguities. We annotated and filtered germline variants through overlap with known SVs (1000G call set, DGV) as well as through overlap with an in-house blacklist of SVs (germline SVs and artifacts called in healthy genomes). As mentioned above, also the split read check helped to remove remaining germline SVs. Finally, we prioritized SVs that were called by more than one tool, or called by only one tool but also confirmed by 1) a CNV changepoint, or 2) at least 3 split reads (in tumor only). Since we found them to be very specific, we also keep Crest-only calls in the high confidence set.

**SV/CNV Annotation.** All predicted copy number and structural variants were annotated with gene overlap (RefSeq, Cancer Census) and potential effect on gene structure (e.g. disruptive, intronic, intergenic). If a predicted SV disrupted two genes and strand orientations are compatible, the SV was annotated as a putative gene fusion candidate. Note that we did not check reading frame at this point. Further annotations include sequence features within breakpoint flanking regions, e.g. mappability, simple repeat content, segmental duplications and Alu repeats.

**SNVs/indels.** The SNV/indel pipeline returned the raw outputs of all variant callers, in VCF format (and for muTect also in TXT format).

We in addition returned the annotated union of all SNVs (*.snv.union.v*.*), union of all indels (*.indel.union.v*.*), and union of all SNVs and indels together (*.union.v*.*), in three formats:

1. VCF - union of individual caller output VCFs, combined using the GATK CombineVariants module;

2. MAF - Mutation Annotation Format, as specified by TCGA (https://wiki.nci.nih.gov/display/TCGA/Mutation+Annotation+Format+(MAF)+Specification) and modified with variant/reference counts columns to be compatible with MSKCC cBioPortal (http://www.cbioportal.org/public-portal);

3. TXT - tab-separated text file, easiest to read and parse but unlike the previous two, this is not a standard, widely accepted file format.

In the *.union.v*.annotated.txt files, the column named “CALLED_BY” indicates the tool(s) that called it. For SNVs this can be:

- mutect
- strelka_snv
- lofreq
- mutect-strelka_snv
- mutect-lofreq
- lofreq-strelka_snv
- mutect-lofreq-strelka_snv

And for indels:

- pindel
- scalpel
- strelka_indel
- pindel-strelka_indel
- scalpel-pindel
- scalpel-pindel-strelka_indel

**SVs/CNVs.** We delivered the raw caller output which comes in a variety of formats (please refer to the individual caller documentation for details). For Delly, these are all files containing “sv.delly”, for BreakDancer “sv.breakdancer”, for Crest “sv.crest” and for NBIC-seq “sv.bicseq”. The output of our SV processing pipeline is in extended BEDPE format (see ReadmeSV.txt) and comes at two levels of confidence:

a. Merged files containing calls from all tools
b. High-confidence files

The full union of calls (a) without any filtering typically still contains many germline variants. The high-confidence variants (b), however, may miss especially low-frequency variants. For an intermediate filtering level we recommend to keep only lines with “known=;” (germline/artifact filter) from the union file.

For more details on files delivered please see the ReadmeSV.txt in the project root directory.

**RNA-Seq Analysis.** The reads were aligned with STAR (version 2.4.4a), and genes annotated in Gencode v18 were quantified with FeatureCounts (v1.4.3-p1). Normalization and differential expression was done with the Bioconductor package DESeq2.

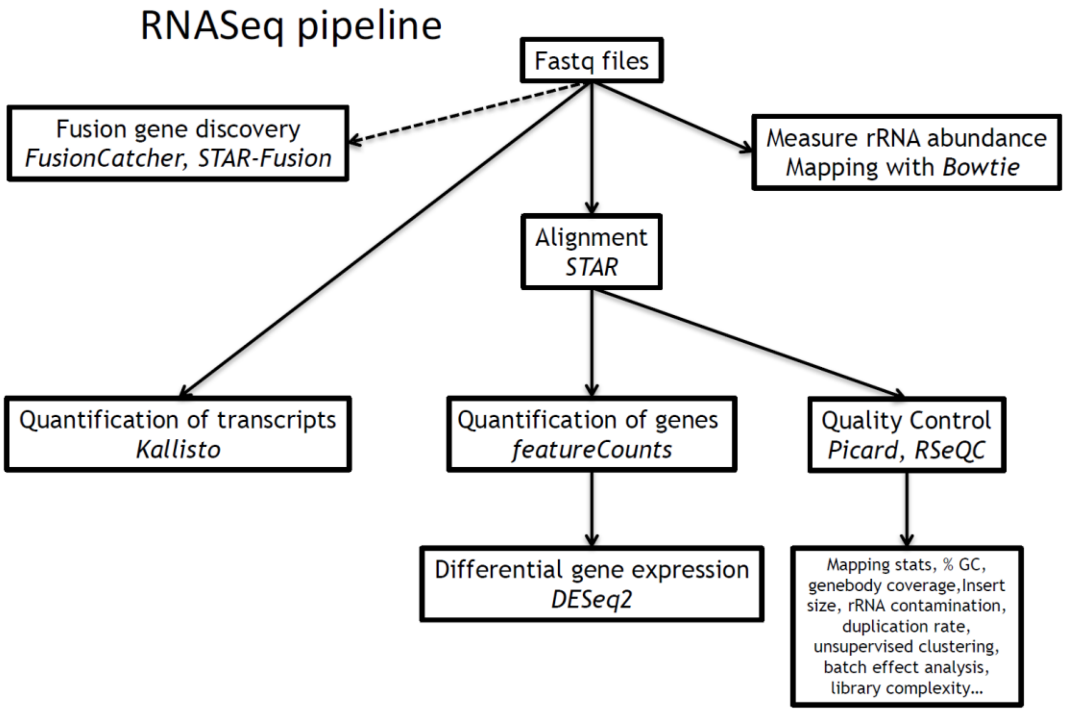


**Methods Figure 7: RNA-Seq Pipeline.**

**Aggregate data analysis.** A master table was generated aligning all the mutation calls across patient samples (https://github.com/kentsisresearchgroup/TargetInductionFailure). The VCF files were processed using Python 3.5.

*Single nucleotide variant (SNV) processing*: SNV VCFs were filtered to retain SNPEFF_IMPACT of "high". "moderate" or "low". The next filter removed any row that did not have one of the following terms as a "SNPEFF_EFFECT":

NON_SYNONYMOUS_CODING, NON_SYNONYMOUS_CODING+SPLICE_SITE_REGION, SPLICE_SITE_ACCEPTOR+INTRON, SPLICE_SITE_DONOR+INTRON, SPLICE_SITE_REGION+INTRON, SPLICE_SITE_REGION+NON_CODING_EXON_VARIANT, SPLICE_SITE_REGION+START_GAINED+UTR_5_PRIME, SPLICE_SITE_REGION+SYNONYMOUS_CODING, SPLICE_SITE_REGION+UTR_3_PRIME, SPLICE_SITE_REGION+UTR_5_PRIME, START_GAINED+UTR_5_PRIME, START_LOST, STOP_GAINED, STOP_GAINED+SPLICE_SITE_REGION, STOP_LOST.

The resulting SNV table was saved and then heatmaps were created with the most recurrent genes mapped to the top of the heatmap. The recurrence was calculated using pivot table from the dataframe.

Variant allele frequencies were calculated as

$$\frac{alternate\ variant\ count}{alternate\ variant\ count + reference\ variant\ count}$$

*Indel processing:* Indel VCFs were filtered to retain SNPEFF_IMPACT of "high" and "moderate". The recurrence was calculated using pivotable from the dataframe. Heatmaps were created with the most recurrent genes mapped to the top of the heatmap.

Additionally all SNVs and indels with variant allele read counts less than 8 were removed.

*Copy Number Variation (CNV) processing:* Since a CNV call could contain multiple genes, each call was multiplied by the number of genes in the variant, creating a row per gene. All the annotations from the CNV was passed on to the new row. This exploding process was applied to all genes in "Cancer_Census=", "DisruptL=" and "DisruptR=", creating one row per gene found in a CNV call. Once done, all rows where the gene was a "neutral" impact were removed. Any row with a missing gene name was also removed.

*Structural Variant (SV) processing:* Since a SV call could contain multiple genes, each call was multiplied by the number of genes in a variant, creating a row per gene. All the annotations from the SV was passed on to the new row. This exploding process was applied to all genes in "Cancer_Census=", "DisruptL=" and "DisruptR=", creating on row per gene found in SV call. Any row with a missing Gene name was also removed.

*Fusion processing:* Fusions were filtered to remove any with the following annotation: banned, bodymap2, cacg, cta_gene, ctb_gene, ctc_gene, ctd_gene, distance1000bp, distance100kbp, distance10kbp, duplicates, ensembl_fully_overlapping, ensembl_partially_overlapping, ensembl_same_strand_overlapping, gtex, hpa, mt, paralogs, readthrough, refseq_fully_overlapping, refseq_partially_overlapping, refseq_same_strand_overlapping, rp11_gene, rp_gene, rrna, short_distance, similar_reads, similar_symbols, ucsc_fully_overlapping, ucsc_partially_overlapping, ucsc_same_strand_overlapping.

We filtered fusions not involving annotated genes and “Fusion_predicted_effect” is not “in-frame.” The final filter removed all “Spanning pairs” that were less than 5, removing lower quality fusion calls.

To accommodate later analysis, we annotated all the genes in the dataframe using the Cancer Gene Census database (version 75).

*Manual verification:* The resulting master table had more than 6000 rows. Approximately 30% were examined in IGV viewer to verify the calls. It became apparent that CNV callers had some false positives. Upon closer examination, we removed the CNV calls from two patient samples. Two samples, PASFLG pre-treatment and PASNKZ pre-treatment, showed amplifications with little SV support. The QC metrics also showed a high autocorrelation which indicates an unevenness of coverage and results in many false focal events being called. Methods Figure 8 illustrates the high copy for the two samples as well as the high QC bias.

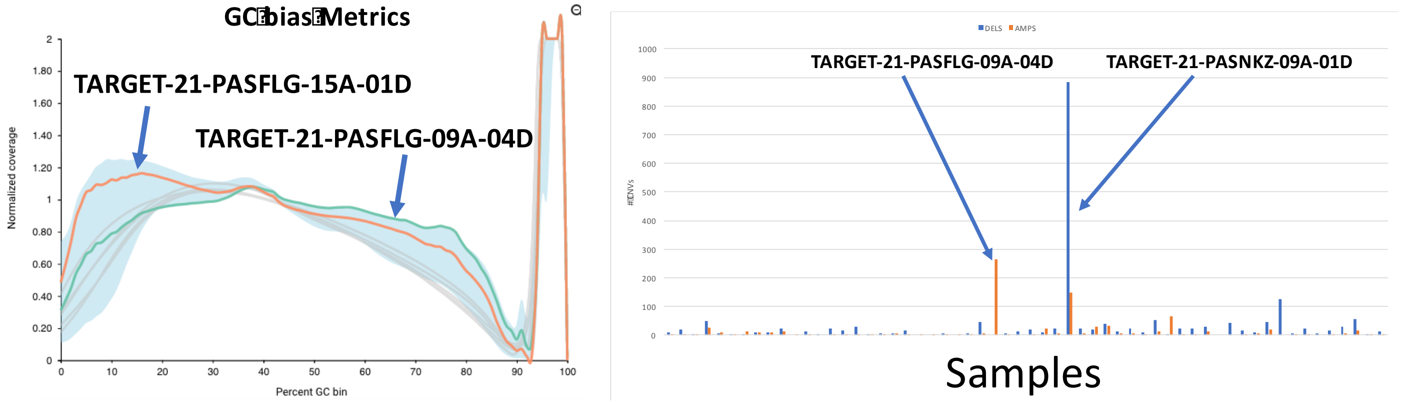


**Methods Figure 8: Two samples show high GC bias.**

The same two samples also show unusually high CNV calls.

Manual verification for the other mutations found that almost all the SNV and indel calls were correct. This is to be expected since the NYGC pipeline required consensus from at least 3 out 5 callers. The few false positives came from regions with very low reads, often near the beginning and end of the chromosome.

*Differential Expression*: Analysis was performed using only the 42 samples with matched pre and post treatment bone marrow. DESeq2 (v1.14.1) was used.

Clustering: Normalized counts were obtained using DESeq2’s counts function with the argument normalized=TRUE. Then, Euclidean distances were calculated on the log2 of these normalized counts using base R’s dist function, and clustering obtained using base R’s hclust function on the Euclidean distances. Base R’s cutree function was used to divide the samples into 4 groups based on the clustering results. This resulted in 3 main groups, plus a fourth group with a single sample that was not used in any downstream analysis. Differential expression was done using DESeq2, running three pairwise comparisons based on the groups obtained by clustering.

PCA: Normalized counts were obtained using DESeq2’s counts function with the argument normalized=TRUE. Then, we took the log2 of these normalized counts to use for unsupervised clustering and PCA. PCA was performed using base R’s prcomp function on the log2 of these counts. Euclidean distances were calculated using base R's dist function, and then clustering obtained using base R's hclust function.

Gene ontology analysis was performed using GOstat, on both upregulated and downregulated genes. These sets of genes were obtained using the union of genes meeting criteria (padj < .001, with the appropriate direction of fold-change) from both pairwise comparisons including group1.

*GSEA*: Differential expression results were obtained using DESeq2 (same version of the program as used to normalize the counts). Three pairwise comparisons were performed over the three groups.

GSEA was performed using GSEA v2.2.1 plus MSigDB v6.0. The hallmark gene set was used. GSEA was run for each set of pairwise comparisons. In the "lfc" results, genes were ranked by their log2FoldChange, with higher magnitude log2FoldChange values weighted higher. Positive log2FC values were assigned to na_pos phenotype and negative to na_neg phenotype. In the spval results, genes were ranked by their adjusted p-value (padj) instead, with lower p-values weighted higher. The sign of the log2FoldChange was still used in these results to determine whether na_pos or na_neg phenotype, but the p-value was used for ranking.

**Hypothesis Testing.** P-values were calculated as indicated in the main text (Fisher's exact, log-rank p, or t-test) using Excel and OriginPro, with corrections for multiple-hypothesis testing.
